# FAIRification of computational models in biology

**DOI:** 10.1101/2025.03.21.644517

**Authors:** Irina Balaur, David P. Nickerson, Danielle Welter, Judith A.H. Wodke, Francois Ancien, Tom Gebhardt, Valentin Grouès, Henning Hermjakob, Matthias König, Nicole Radde, Adrien Rougny, Reinhard Schneider, Rahuman S. Malik-Sheriff, Kirubel Biruk Shiferaw, Melanie Stefan, Venkata Satagopam, Dagmar Waltemath

**Affiliations:** Luxembourg Centre for Systems Biomedicine (LCSB), University of Luxembourg, Luxembourg; Auckland Bioengineering Institute, University of Auckland, New Zealand; Luxembourg National Data Service (LNDS), Luxembourg; Medical Informatics Laboratory, University Medicine Greifswald, Germany; European Molecular Biology Laboratory, European Bioinformatics Institute (EMBL-EBI), UK; Institute for Biology, Institute for Theoretical Biology, Humboldt University of Berlin, Germany; Institute for Stochastics and Applications, University Stuttgart, Germany; Medical School Berlin, Berlin, Germany

## Abstract

Computational models are essential for studying complex systems which, particularly in clinical settings, need to be quality-approved and transparent. To enhance the communication of a model’s features and capabilities, we propose an adaptation of the Findability, Accessibility, Interoperability and Reusability (FAIR) indicators published by the Research Data Alliance to assess models encoded in domain-specific standards, such as those established by COMBINE. The assessments guide FAIRification and add value to models.

## Main

### Introduction and background

Computational models are useful tools for studying complex systems^1^. Data security and patient safety require these models to be particularly quality-approved when used in a biomedical context^2^. Reasons for the low uptake of computational modelling in clinical practice include limited transparency of the model code, lack of standardisation of the settings indicating how the model can be simulated, insufficiently clear specification of the model parameters, and low reproducibility of simulation results^3,4^ as well as a lack of credibility of the models^5^. One step towards improving credibility and reproducibility of scientific results in general is the commitment to adhere to the principles of Findability, Accessibility, Interoperability and Reusability (FAIR)^6^.

A FAIR assessment is a structured method to determine the extent to which a data object conforms to the FAIR guiding principles^6^. A common set of domain-agnostic FAIR indicators has been published by the Research Data Alliance (RDA). Subsequently, the IMI FAIRplus project (https://fairplus-project.eu/, accessed February 2025) provided a template for FAIR assessment of data and metadata distinguishing three levels of agreement (0: no information/not fulfilled; 1: information fulfilled; and NA: not applicable)^7^. Importantly, metadata is a key component for every FAIRification journey. Missing metadata reduces the overall FAIRness level of the resources examined (see the FCB068 recipe^8^ from the FAIR Cookbook^9^). The results of a FAIR assessment, often visualised as so-called radial column charts, indicate the strengths and weaknesses of the FAIRness of a dataset. An analysis of the radial columns leads to a FAIRification plan which guides the steps to improve FAIRness.

A low initial score in a baseline FAIR assessment is the rule and not an exception. At the same time, the complete fulfilment of all FAIR indicators requires substantial effort and expert knowledge. In most cases, a 100% FAIRness score is not desirable. Depending on the specific application area, a subset of RDA FAIR indicators might not be essential to provide good data objects. It is more important to identify the necessary indicators in a specific research context and then work towards a higher FAIRness score of the respective FAIR principles. It is important to note that many researchers today are aware of the FAIR principles. In most cases, a substantially higher FAIR score is easily reached if the already available information is structured and formatted correctly with support of FAIR experts. Repositories usually encourage and actively support FAIRness by providing standards and guidelines for representing scientific results. While the final FAIRness of a model is sole responsibility of the developers, model repositories can define the minimal FAIRness required to list a model by explicitly requesting FAIR indicator related information during the model submission process. The amount of information requested varies between repositories as an increasing complexity of the submission process and is also a putative obstacle preventing for example less experienced researchers from using a repository at all. Many model repositories were set up before the FAIR principles were articulated and can be considered pioneers for transparency and reproducibility in scientific modelling. Examples include the BioModels^10^ database (www.ebi.ac.uk/biomodels/, accessed February 2025) and the Physiome Model Repository^11^ (PMR, https://models.physiomeproject.org, accessed February 2025).

In 2022, we launched the FAIR COMBINE Archive Indicators project (https://fair-ca-indicators.github.io/, accessed February 2025) as a community effort for the FAIR assessment of models and simulation studies within the COmputational MOdeling in BIology NEtwork (COMBINE), the largest scientific community for systems biology modelling^12^. Building on the work of the RDA, the FAIRplus project and our own previous work on the FAIR assessment of biomedical data objects^13^, we adapted the RDA FAIR indicators template provided by the FAIRplus project to cover the features of the COMBINE simulations. We assessed both the computational models themselves and the COMBINE archives, which represent the set of files needed to reproduce a virtual experiment^14^. Figure 1 summarises the workflow of the FAIR COMBINE Archive Indicators project, which was conducted under the umbrella of the RDA/European Open Science Cloud (EOSC) Future project (no. 101017536).

**Figure 1:**
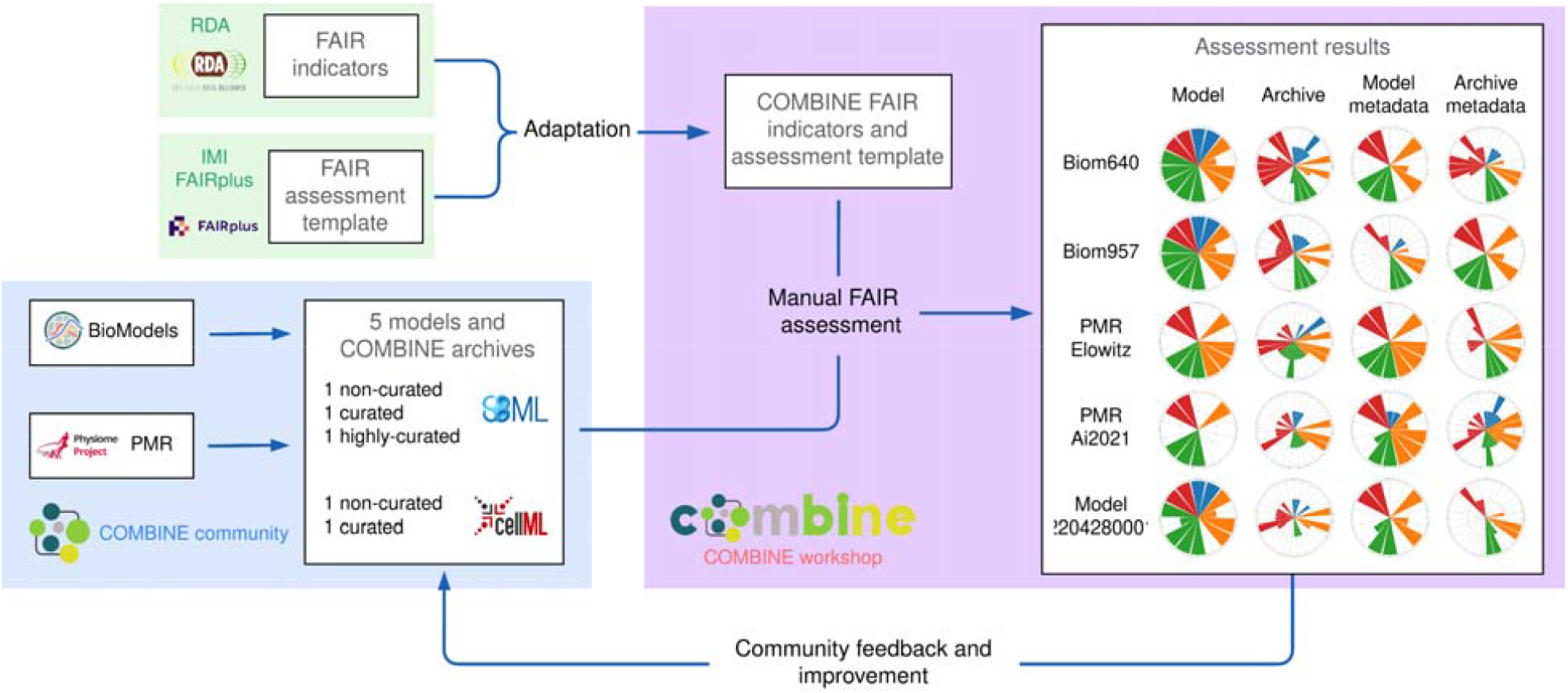
Workflow of the FAIR COMBINE Archive Indicators project: an iterative and integrative community-level process to adapt the RDA FAIR indicators to the specificities of COMBINE models and archives. We extended the FAIRplus template of RDA FAIR indicators and performed FAIR assessment for 5 types of COMBINE models and archives from major model repositories (BioModels and PMR) at different curation stages.

### Methods

We focused on assessing the FAIRness of COMBINE models and archives. Publicly available models encoded in the Systems Biology Markup Language (SBML)^15^ and CellML^16^ were selected from BioModels and PMR and assessed at different stages of model curation. The FAIR assessment of COMBINE archives included:

- the omex^14^ standard format for the archive itself.
- SBML or CellML files stored inside the archive.
- the Systems Biology Graphical Notation (SBGN)^17^ files for graphical representations of the network.
- the Simulation Experiment Description Markup Language (SED-ML)^18^ files for standardised descriptions of simulation setups.

The COMBINE FAIR indicators were defined in a multi-step community process during two FAIR-dedicated workshops running at the main COMBINE conference and the corresponding hackathon in October 2022 and April 2023. These workshops were preceded by two other COMBINE workshops, held in October 2021 and April 2022, where we initiated the discussion of the FAIR indicators. We also organised a series of five online open hackathons to revise and finalise the assessment (February - March 2023). We articulated a community-consensus to adapt the FAIR indicators to both the COMBINE models (which only capture the model formulation) and the archives (which include supporting resources for the model). Details on the approaches and project activities are provided in the Online Methods sections.

### Results

The final version of the COMBINE FAIR indicators distinguishes 42 indicators for models and 42 indicators for archives^19^. We evaluated the indicators on five different models and associated archives from BioModels and PMR at different stages of curation (Table 1). Details on the complete FAIR assessment and visualisation of the FAIRness score for each of these models are given in SupplementaryFile_1 and SupplementaryFile_2, respectively. As we agreed to acknowledge any effort towards FAIRness, we marked several exceptional cases with the score of 0.5 (i.e. between 0 and 1), where considerable work had already been done but the indicator is not yet fully satisfied. In general, the FAIR assessment depends on the evaluation context and is characteristic to the resource itself. The FAIR score indicates how the essential and non-essential FAIR indicators, respectively, are fulfilled by the assessed resource. Also, the score should be interpreted in the context of the same resource over time as improvements are made; the FAIR score is not informative for comparing between resources. Moreover, rather than focusing on the FAIR scores, the FAIR assessment of multiple resources can identify common gaps impacting the FAIRness among the resources. Considering this, in our assessments our aim was to identify key components (specific to each evaluated category) that can be further improved from the FAIR point, with the ultimate goal to increase the reproducibility of the resources.

**Table 1:**
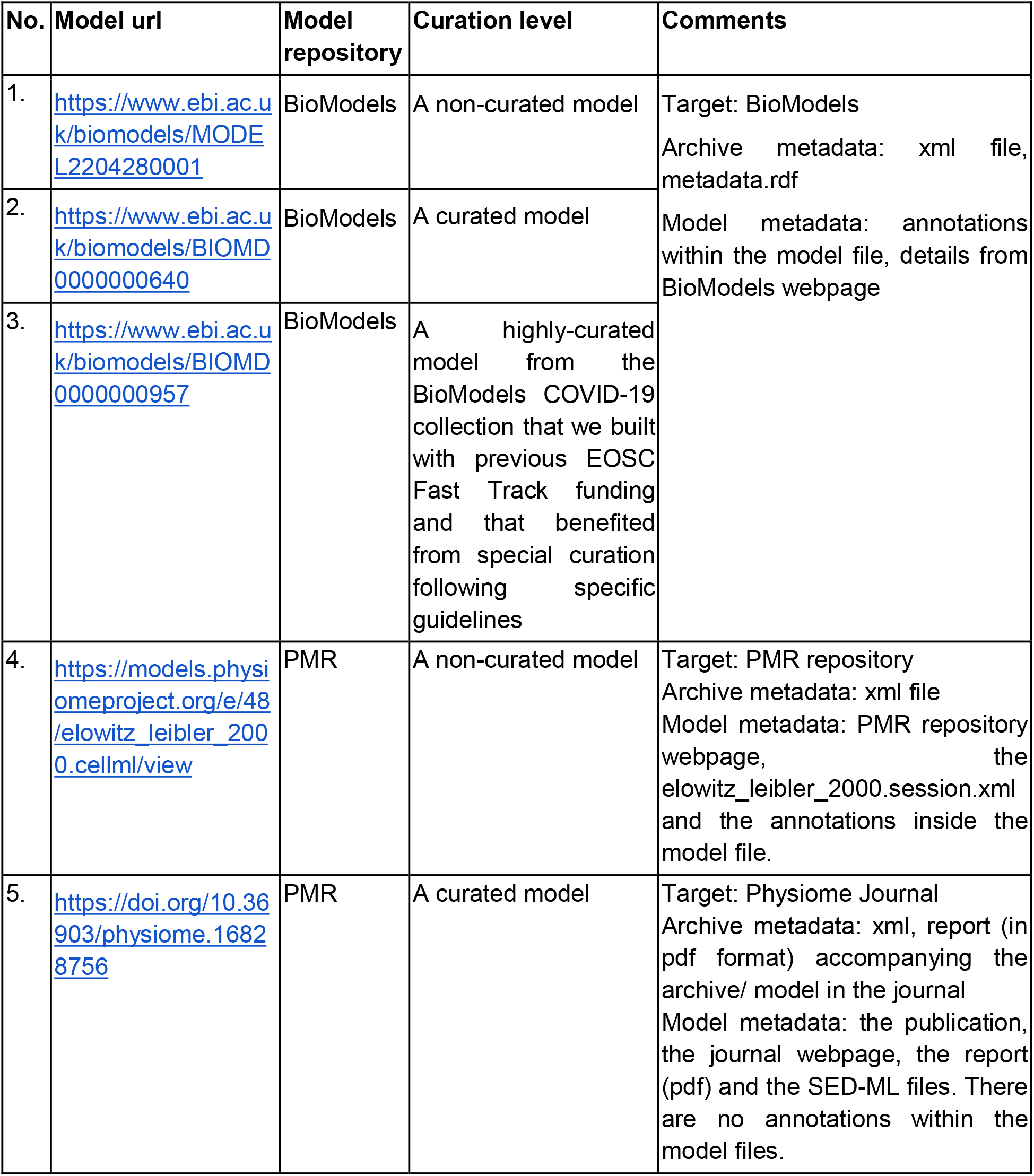
Details of the 5 selected categories of the models and archives (considering the curation level) used to perform the FAIR assessment using the COMBINE-adapted RDA FAIR indicators. The metadata of archive/ model represent key FAIR assessment components. We give details on metadata composition considered for each category specifically.

The results of our FAIR assessments confirm that the COMBINE community has already achieved a reasonable FAIRness level. In Figure 2, we show as an example the FAIR assessment results for one of the models used during the workshops and hackathons (https://models.physiomeproject.org/e/48/elowitz_leibler_2000.cellml, accessed February 2025). The result illustration is made using the FAIR-Viz^20^ tool ((https://faircombine.streamlit.app/). First, the use of COMBINE standards contributes to the interoperability and reusability at the community level. In the evaluations, the corresponding indicators, such as the Reusability indicator on the model using community standards (CA-RDA-R1.3-01Model), were scored as full-filled (score = 1) because the models used the SBML and CellML standard formats. Also, the use of the repositories (BioModels and PMR) contributed to the FAIRness of the assessed resources. For example, the BioModels and PMR platforms both provide API and access protocol for accessing the models and archives, fulfilling the Accessibility indicators on automatic access and on free access protocol (CA-RDA-A1-05Model and CA-RDA-A1.1-01Model, score = 1).

**Figure 2:**
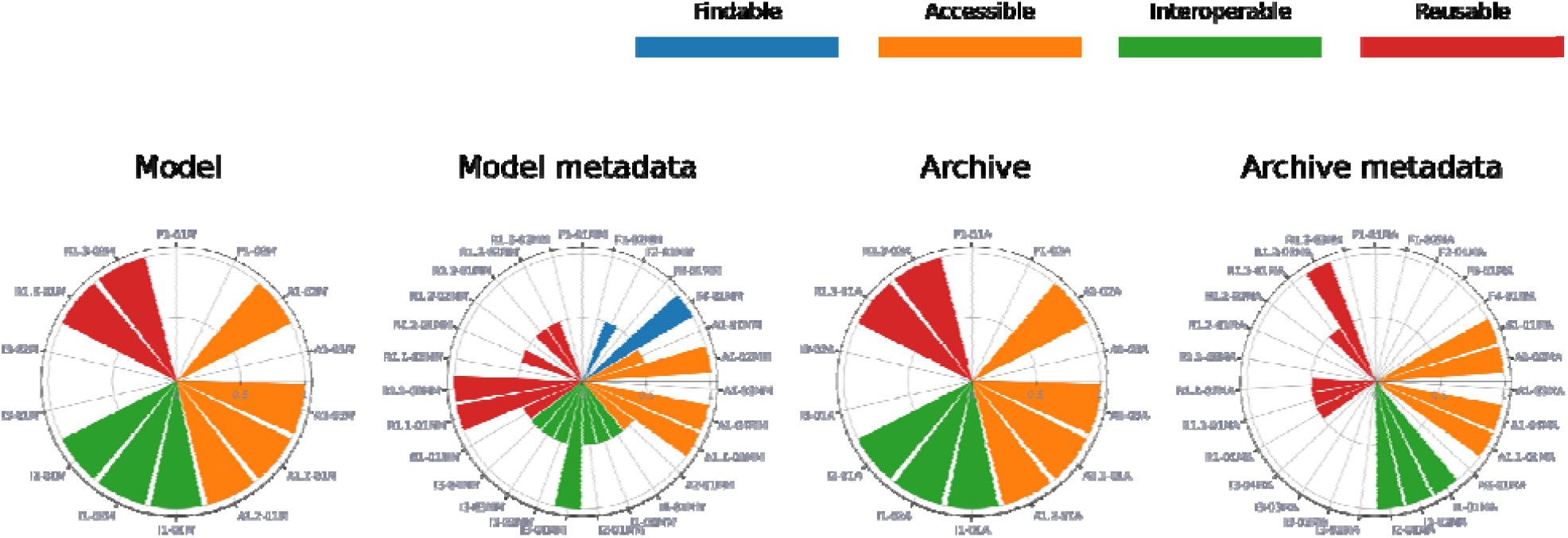
Example: FAIR assessment results for one of the model types considered during the workshops and hackathons - https://models.physiomeproject.org/e/48/elowitz_leibler_2000.cellml. Results are shown for the resource itself (Model or Archive) and the metadata component (namely Model metadata or Archive metadata), for each FAIR indicator. The result illustration is made using the FAIR-Viz^20^ tool. Legend: Findability (blue), Accessibility (orange), Interoperability (green), Reusability (red).

The FAIR assessment has also identified potential for further FAIRification of models and archives. Metadata is a key component of the FAIR principles, and the absence or limited availability of metadata highly affects the FAIRness of data objects, regardless of the level of curation of the selected resources. Within the context of this work, metadata was partially available, but it was often distributed across several locations (including the xml file, the corresponding webpage or publication, within the models themselves) rather than being accessible in a single location, making it difficult to access information about the resources, especially in an automatic manner.

### Discussion

A FAIR assessment before publishing a simulation study provides details on the reusability of the contained data objects. It is essential for sustainable modelling to provide all relevant metadata and that the work is licensed and accessible according to the chosen licenses. As a maintainer of a model repository or simulation tool, it is advisable to perform a FAIR assessment as part of the submission process to ensure a high quality of published research assets.

Publishing computational models in repositories alongside the journal publication has been established as part of good scientific practice within the COMBINE community and is encouraged by many journals. Upon publication of reusable simulation studies in open model repositories certain metadata should be provided by the submitting authors and the model curators to ensure reproducibility of the scientific results. The BioModels team, for example, has developed a distinct checklist to address the high rate of irreproducible simulation studies^3^. The SBGN community has provided “quick tips for creating effective” and meaningful visual model representations^21,22^.

A structured and coordinated approach to assess the actual FAIRness of COMBINE resources did not exist yet. Despite all the effort taken in different projects, a single assessment protocol was missing. With the help of an EOSC fund, a work group at COMBINE therefore adapted the RDA FAIR indicators matrix^7^ and optimised it for the assessment of both computational models and simulation studies (archives) within the ecosystem of COMBINE. Many community members, despite having a high interest in the FAIR principles, are not themselves experts in FAIR assessment tools and FAIRification workflows. The workshops and expert discussions with developers of computational models, modelling tools, and infrastructure for code sharing led to COMBINE FAIR indicators^19^. Based on what has been defined as the necessary FAIR indicators and our lessons learnt from the FAIR assessments (Table 1), we formulate the following recommendations for everyone wishing to provide FAIR computational models:

1. **Metadata should be structured and standardised, should be provided in a separate file, and should remain available even if the computational resources are deleted**. Currently, models and metadata are often mixed within the model encoding. For example, SBML core contains model elements and metadata. We propose to strictly separate data and metadata, which means to provide separate metadata files for models in line with the recommendation for the harmonisation of metadata by Neal et al^23^. In addition, the metadata needs to be both structured (to be machine-readable) and standardised (according to available community standards^23,24^). Separate provision of all metadata about an archive will enable users to explore and compare models and archives directly, without having to first parse the full file set to identify and extract specific parts related to metadata for further inspection. In addition, the metadata should remain available even if the resources have been deleted.
2. **Quality of metadata should be assessed in addition to its pure existence**. Data objects with rich metadata may still contribute little to the actual understanding of the data if, for example, the metadata is not domain-specific, is not correct, or is too general. FAIR metadata does not itself guarantee that the metadata is informative, i.e. has a high information content^25^.
3. **Basic provenance should be recorded**. Changes to models, archives and metadata files should be documented using provenance standards. It also needs to be discussed how to handle the metadata when a data object (model or archive) is deleted. We consider the metadata of simulation studies particularly essential for understanding and reusing the model as it often encodes details that are needed to calibrate and execute the model code, e.g. simulation algorithms and initial parametrisation^24,26^. Therefore, we argue that such metadata needs to be tracked using provenance information to make the longitudinal history of simulation setups available.
4. **Each relevant component of a simulation study should have a unique ID**. As a consensus of our working group, we believe that a crucial next step for FAIRification of models and archives is the consequent use of IDs (indicator CA-RDA-A1-01). Each model, model version, metadata file, and the archive itself should have their own IDs to ensure a consistent reference point, to track changes between versions, to refer to individual model versions, and to be able to put files into relation, even across different archives.

The COMBINE community is committed to open and transparent science. In line with this mission, we would like to encourage researchers to publish their FAIR baseline scores, so that the scientific community is not hampered by a feeling of unfairness, when scoring low in FAIRness. Reaching a high FAIRness score (and everything above 80% is high) is a team effort, and requesting support from FAIR experts is highly recommendable.

Throughout the project we have disseminated our working strategies, results, lessons learned and challenges to the broader Open Science community, and we always emphasised on the benefits of following FAIR principles for the reproducibility of computational models.

In order to facilitate FAIR assessments, we also worked on the development of a semi-automatic FAIR assessment tool for COMBINE models and archives, where some RDA FAIR indicators are automatically assessed based on the properties of the inspected resources (models or archives). For example, the tool automatically assesses features with key impact on the FAIR indicators, such as the existence of metadata files. Other FAIR indicators should be evaluated manually with user input. The FAIR assessment tool can help modelers and curators increase the FAIRness of their own research and identify the weak points with respect to model reusability and result reproducibility. The tool is openly available as a prototype (backend: https://github.com/FAIR-CA-indicators/fair-ca-indicators-backend, accessed February 2025; frontend: https://github.com/FAIR-CA-indicators/fair-ca-indicators-frontend, accessed February 2025) under Apache License 2.0.

Finally, we plan to use the presented list of COMBINE-adapted RDA FAIR indicators to formulate community guidelines for the FAIRification of models and archives during development and publishing. One immediate step is to encourage the community to provide metadata separated from the actual model and archive, as detailed as possible to facilitate the usability and reusability of the work. We believe that the COMBINE FAIR project is the first step towards systematic FAIR assessment of computational models, leading to more FAIRness and thus faster curation and better reproducibility of computational biology studies.

## Online Methods

In September 2022, we launched the FAIR COMBINE Archive Indicators project under the umbrella of the RDA/EOSC Future project (https://fair-ca-indicators.github.io/, accessed February 2025) to adapt the FAIR indicators^6^ to computational biology models and archives of simulation studies^12^. Together with the COMBINE community, we adapted the RDA FAIR Data Maturity Model^7^ indicators, following the template provided in the IMI FAIRplus project (https://fairplus-project.eu/, accessed February 2025), to the COMBINE specifics. COMBINE models and archives are available for the broader research community in open model repositories such as BioModels^10^ and Physiome Model Repository (PMR)^11^. Besides the model encoding in the SBML^15^ or CellML^16^ formats, a standardised graphical representation (SBGN^17^) and a recipe for reproducible experiments (SED-ML^18^) are typically provided. The COMBINE archive (omex^14^ format) contains all data necessary to reproduce a simulation experiment.

### FAIR-related work prior to the FAIR COMBINE archive project

Prior to the FAIR COMBINE Archive Indicators project, we performed the FAIRness assessment using the RDA FAIR indicators in several biomedical projects^13^, including the IMI BIOMAP (https://biomap-imi.eu/, accessed February 2025) and the EU ERAPerMed HeartMed (https://heartmed.pages.uni.lu/, accessed February 2025) projects. In addition, we were participants of the IMI FAIRplus project. We also coordinated the FAIRness assessment^27^ using the RDA indicators for the biological diagrams available in the Molecular Interaction NEtwoRk VisuAlization platform (https://minerva.pages.uni.lu, accessed February 2025) for the Parkinson’s disease map (https://pdmap.uni.lu, accessed February 2025) and the COVID-19 Disease Map (https://covid.pages.uni.lu/, accessed February 2025), within the Disease Maps Project (https://disease-maps.io/, accessed February 2025), which concentrates on the development of computable and comprehensive knowledge repositories for translational medicine research. Most of the FAIR principles were assessed^27^, including the provenance of source information, the use of stable annotations for the map components, and the reusability of the content due to the use of the Systems Biology standards.

Within the COMBINE community, we organised two FAIR-dedicated workshops in October 2021 and April 2022, where we conducted a series of introductory presentations and discussions about the FAIR principles, in general, and about the use of RDA FAIR indicators, in particular, with regard to the FAIR assessment of the COMBINE computational models. We also discussed with the community about benefits and issues of performing the FAIR assessment with respect to model quality. We analysed the set of FAIR indicators from the RDA Data Maturity Model^7^ and concluded that they can be used in the context of COMBINE, but some degree of adaptation was needed to address the specifics of the COMBINE archives.

### Community workshops for adaptation of RDA FAIR indicators to COMBINE

We organised a series of community workshops and meetings at which we asked for community contribution, dissemination, and consensus during the major phases of the project, including i) adapting the RDA FAIR indicators to the COMBINE models and archives, ii) performing the FAIR assessment for several types of computational resources considering their curation level, and iii) discussing further needs and steps, including the development of community guidelines for FAIRification of the COMBINE simulations as well as of a tool to facilitate the FAIR assessment by completing several indicators automatically, given the input settings of the models and archives.

#### COMBINE 2022

At the annual community meet-up, COMBINE 2022^28^, we organised a workshop specifically for the FAIRness assessment of COMBINE archives. We aimed to raise awareness for FAIR in the COMBINE community and to specifically promote the assessment of the quality of research outcomes using a coordinated FAIR assessment. Circa 200 community members were present at the COMBINE 2022 meeting and were informed about the existence and possible use of FAIR assessment tools, specifically the RDA indicators. The workshop itself attracted 30 participants (developers and curators of COMBINE models, experts on the BioModels and PMR open model repositories) who discussed the first version of RDA indicators for FAIR assessment of COMBINE archives. Participants worked in five groups, each group focusing on the assessment of a specific COMBINE model and archive, as follows (all the urls from this list were accessed and verified on February 2025):

1. a non-curated model from BioModels: https://www.ebi.ac.uk/biomodels/MODEL2204280001;
2. a curated model from BioModels: https://www.ebi.ac.uk/biomodels/BIOMD0000000640;
3. a highly-curated model from the BioModels COVID-19 collection that we built with previous EOSC Fast Track funding and that benefited from special curation following specific guidelines: https://www.ebi.ac.uk/biomodels/BIOMD0000000957;
4. a non-curated model from PMR: https://models.physiomeproject.org/e/48/elowitz_leibler_2000.cellml/view;
5. a curated model from PMR: https://doi.org/10.36903/physiome.16828756.

During this workshop, participants decided to perform the FAIR assessment on the COMBINE archive instead of assessing FAIRness for the COMBINE model itself (which captures the model formulation only). We continued working with FAIR indicators as available in the RDA FAIR Data Maturity Model^7^, and evaluated the applicability of every indicator for the COMBINE archive. The workshop participants also explored the priority levels of the indicators.

Participants familiarised themselves with the RDA FAIR indicators and the provided FAIRplus template in smaller groups. Not all groups were able to finalise the assessment, as for most participants, this was the first time they had been exposed to the RDA FAIR indicators. The participants worked consciously and practically on the indicators. An important learning goal was the understanding that the model building and publication processes benefit from following and applying the FAIR indicators from the model development stage onwards, not only after the model is released. One key open question, which was addressed and decided upon in the later workshops, was whether to perform the FAIR assessment at the level of individual models (computational formulation file only), or archives (aggregated resources including the model itself and supporting files) or both in order to maximise the benefits of the FAIR assessment across the entire COMBINE community.

#### Online hackathons

While the previous workshops were well-perceived by the community and received much attention in the COMBINE community at large, it became obvious that a detailed FAIR evaluation of the COMBINE archive demands much more work on the conceptual and implementation level. Importantly, there was a community-consensus that having the FAIR assessment for one category only (archives or models) would still miss specifics of the other category. Thus, following several bi-weekly community meetings, we decided to systematically perform the FAIR assessment targeting both the COMBINE models and archives. We organised a series of five online hackathons to revise and finalise the assessment of the five example models. Hackathons were open to the community and accompanied by an open call for feedback via github issues (see details at https://github.com/FAIR-CA-indicators/CA-RDA-Indicators/issues, accessed February 2025). Each session of the hackathon was a moderated one-by-one manual FAIR assessment of the chosen model and archive, respectively, based on the latest version of indicators. The working groups included participants with expertise on FAIR, COMBINE as well as on the BioModels and PMR model repositories.

The applied FAIRplus template includes 41 RDA FAIR indicators with 3 levels of priority (namely Essential, Important and Useful) and a binary scoring system: 1 meaning that a specific indicator was fulfilled, 0 otherwise. If the indicator is not specific to the given context, it is marked as not-applicable (NA). The FAIRplus template also distinguishes between Data and Metadata as key components. In our work, having decided to perform assessment for both models and archives, we first mapped the FAIR components to the COMBINE resources as follows. The models and the archives (within all the inner supporting files) were considered as Data in the FAIR template, and the metadata of the models and of the archives as Metadata. Consequently, the Data-related indicators were renamed to Model indicators and to Archive indicators. Similarly, the Metadata-related indicators were renamed to Model Metadata indicators and to Archive Metadata indicators. This extended the initial set of 41 RDA FAIR indicators (focusing on Data and Metadata, in general) to a new set of 82 indicators focusing on the Model and its metadata (41 indicators) and the Archive and its metadata (41). We anticipated the need for metadata to be in compliance with a machine-understandable cross-community standard in order to permit automatic retrieval and interrogation of the COMBINE resources (based on their metadata) from outside of the COMBINE community. Consequently, we proposed the introduction of 2 new Reusability indicators on the metadata of the archive and of the model, respectively, to comply with cross-community standard, namely: “CA-RDA-R1.3-03MA: Metadata of archive is expressed in compliance with a machine-understandable cross-community standard” and “CA-RDA-R1.3-03MM: Metadata of the model is expressed in compliance with a machine-understandable cross-community standard”. Thus in total, we defined a set of 84 RDA FAIR indicators adapted to COMBINE. Further, we discussed and finalised the priority and the description of each indicator.

Although the FAIRplus template uses binary scores (0 or 1) for indicator assessment, during the 5 sessions of the FAIR assessment, we agreed to allow the intermediate score of 0.5 in exceptional cases to acknowledge existing information even if not completely satisfying the indicator, e.g. if the requested information is provided in a different form than the one expected by the indicator’s description. For example, in the assessment of the PMR curated model and archive, the Reusability indicator CA-RDA-R1.2-01MM (“Metadata of model includes provenance information according to community-specific standards”) was scored value 0.5, with the explanation that some standardised metadata information was available in the PMR Journal and in the repository itself. However, the metadata is not available in separate files. Therefore we chose this partial scoring strategy to encourage the COMBINE community to apply the FAIR principles, and we anticipate that some of the current cases will be solved once the community evolves on following the FAIR indicators during the model development, release and curation. We also plan to add this partial scoring situation in our future work, perhaps discussing and learning from other ongoing FAIR initiatives on software.

During our working FAIR assessment sessions, we also noticed a level of dependency between indicators, where the assessment of the former ones as non-fulfilled would automatically determine the assessment of specific latter ones to non-fulfilled. For example, if an archive does not include any references to other data, it is evident that the archive will not include qualified references either. Thus, if the CA-RDA-I3-01Archive indicator on “Archive includes references to other data” is assessed to non-fulfilled, with score = 0, then the CA-RDA-I3-02Archive indicator on “Archive includes qualified references to other data” can be automatically assessed as non-fulfilled, also with score = 0. Furthermore, we observed that some FAIR indicators are supported by the model repositories and can also be assessed automatically. For example, the BioModels repository has an overall licence that must be agreed to by all inner models. Therefore, the RDA indicators on the licence (e.g. CA-RDA-R1.1-01MM) can be completed automatically for models stored in BioModels. We discussed the next steps, including the development of a semi-automated FAIR assessment tool to support the community in the FAIR assessment process, mainly by (1) automating the assessment for a subset of related indicators and (2) providing a visual interface to the assessment results and the suggested improvements.

#### HARMONY 2023

We continued work on the adaptation of the RDA FAIR indicators to the COMBINE simulations and we introduced a prototype of the semi-automatic evaluation tool. The development of the tool integrated the feedback from the community and continued until the end of the project.

### Alignment with the European Open Science Cloud (EOSC) and the Research Data Alliance (RDA) initiatives

A key challenge for EOSC is to develop broadly applicable guidelines, tools, and workflows and to support their cost-effective specialisation for community-specific needs^29^. The application of the broad RDA FAIR indicators to COMBINE standards is a prime example for this strategy. Our main challenge was achieving a community-level accepted set of FAIR indicators to capture the specifics of the COMBINE resources. For this, we relied on the community members being proactive. We continuously involved the community by organising workshops, communicating transparently and acknowledging involvement, or via online dissemination of preliminary results. Moreover, we introduced our project goals and discussed our strategies and results in several scientific conferences and meetings in the broader Computational Biology and Open Science community (including the ICSB 2022, EOSC Symposium 2022, FAIRDOMHub User Meeting 2023, 12th EURINT INTERNATIONAL CONFERENCE 2023, 1st Conference on Research Data Infrastructure 2023, ISMB/ECCB2023). We made connections to other relevant initiatives and projects such as FAIR Principles for Research Software (FAIR4RS Principles: https://www.rd-alliance.org/group/fair-research-software-fair4rs-wg/outcomes/fair-principles-research-software-fair4rs, accessed February 2025), NFDI4DataScience (NFDI4DS: https://www.nfdi4datascience.de/, accessed February 2025), NFDI4Health (https://www.nfdi4health.de/, accessed February 2025), Network University Medicine Germany, etc. All our work components were open and freely-accessible and connected via our project website (https://fair-ca-indicators.github.io/, accessed February 2025) to zenodo (dissemination material^30–33^) and github for code files. Our aim is to foster the connection between the EOSC/ RDA FAIR and COMBINE communities as well as to contribute to raising the potential application of FAIR indicators in other domains towards Open Science.

With this project, the interaction between RDA and COMBINE communities were strengthened. The COMBINE community benefits from reusing the existing RDA FAIR indicators. The RDA became more prominent in the COMBINE community.

## Supporting information

Details on the complete FAIR assessment and visualisation of the FAIRness score for each of these models are given in SupplementaryFile_1 and Suppleme

the complete FAIR assessment and visualisation of the FAIRness score for each of these models are given in SupplementaryFile_1 and SupplementaryFile_2

## Acknowledgement

We thank all participants of the COMBINE community for the work on adapting the FAIR indicators to the COMBINE models and archives.

