## Supplementary material for "FAIRification of computational models in biology": the complete FAIR assessment and visualisation of the FAIRness score for each of these models are given in SupplementaryFile_1 and SupplementaryFile_2

### **SupplementaryFile_2: Visualization of FAIR assessment results for all models included in Table 1**

The result illustration is made using the FAIR-Viz tool (<https://faircombine.streamlit.app/>, github: <https://github.com/matthiaskoenig/fair-ca-visualization>). Legend: Findability (blue), Accessibility (orange), Interoperability (green), Reusability (red).

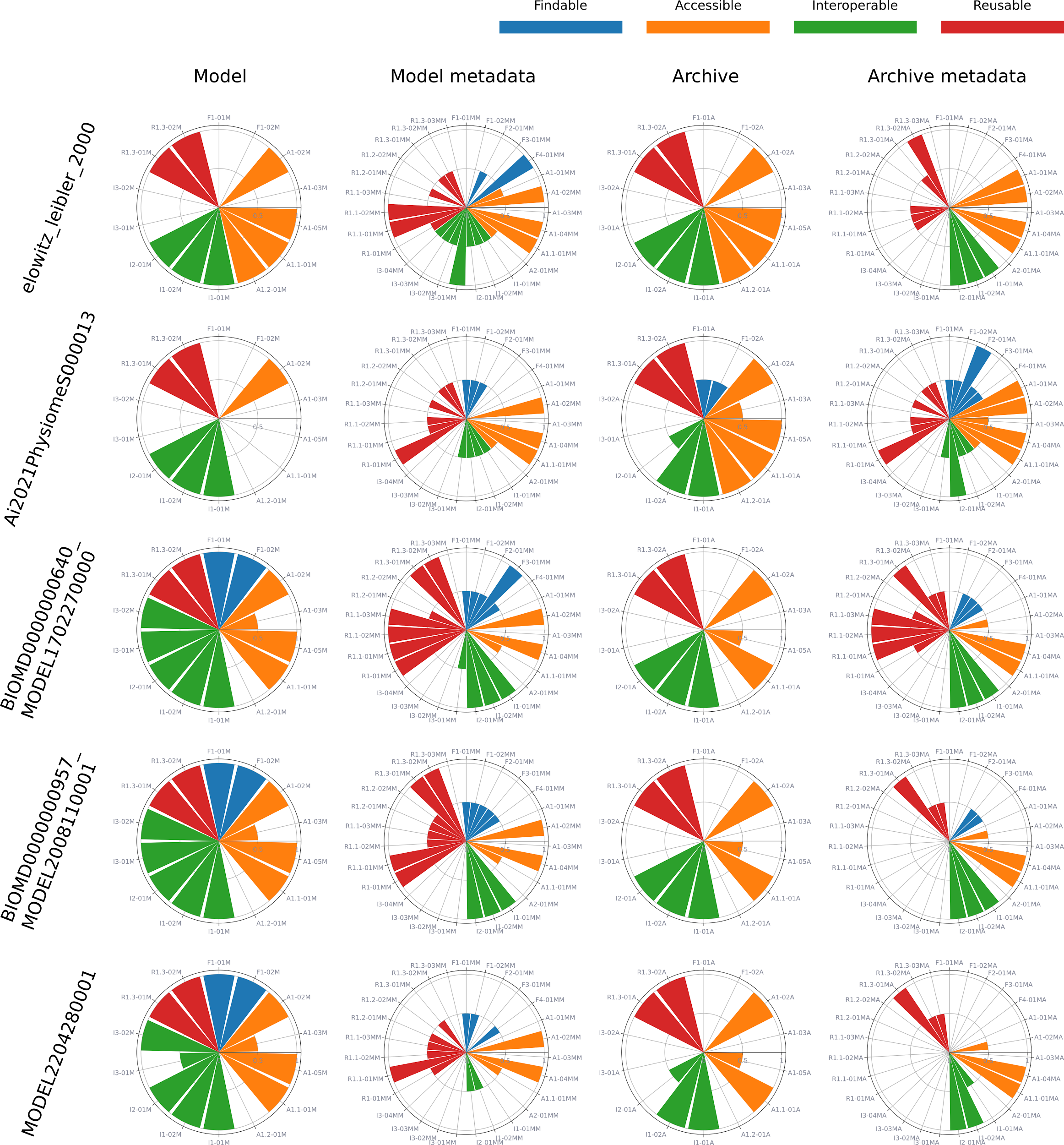
